# Texas 2-Step: A new Model for YcgR::c-di-GMP Action at the Flagellar Motor

**DOI:** 10.1101/2025.08.21.671521

**Authors:** Nabin Bhattarai, Wangbiao Guo, Jonathan D Partridge, Rasika M. Harshey

## Abstract

YcgR is a c-di-GMP effector that inhibits chemotaxis and swimming speed in *Escherichia coli* and *Salmonella*. Genetic, biochemical, and structural studies suggest that YcgR interacts with both the bidirectional flagellar rotor and the stator to bias rotation toward counterclockwise (CCW) and reduce motor speed, but the underlying mechanism remains unresolved. Recent cryo-EM structures revealing conformational changes in the rotor–stator complex during directional switching suggested to us a mechanism by which YcgR acts. We call this the **Texas 2-Step** model, after the country dance in which partners move smoothly in a CCW arc with quick steps followed by slow ones. In this model, YcgR first binds a MotA subunit when the rotor adopts the CCW conformation, in which stators are largely displaced from the C-ring. In the next step, the rotating MotA pentamer delivers YcgR to the rotor protein FliG, thereby slowing motor speed. We provide evidence for the first step of this model, offering testable predictions for future work.

**Importance:** The mechanism of YcgR action has been investigated by multiple laboratories using diverse approaches, yet no consensus has emerged. Some studies implicate the rotor, others the stator. A key complication is the involvement of four interacting proteins—MotA, FliG, FliM, and YcgR—with multiple contact sites in several of them. Recent rotor–stator cryo-EM structures revealing conformational changes during directional switching suggested a mechanism that we set out to test. Our experiments show that rotor conformation is crucial for YcgR function.

## Introduction

Among the many outputs of c-di-GMP signaling, a ubiquitous one is inhibition of bacterial motility (1). The inhibitory effect on flagellar motility is well-studied in *E. coli* and *Salmonella* and is orchestrated via interaction of the c-di-GMP effector YcgR with both the flagellar rotor and stator (2–8). In these bacteria, YcgR expression is under the control of the flagellar regulon, where a three-tiered regulatory cascade integrates various environmental cues and ensures an ordered assembly of the flagellum (9). Curiously, both YcgR and PdeH (also called YhjH), a strong phosphodiesterase that degrades c-di-GMP, are expressed in the last tier of the cascade (10–13). PdeH is expected to keep c-di-GMP levels low when cells are all set to move, while YcgR is likely on guard to arrest motility if environmental conditions elevate c-di-GMP levels via the multiple diguanylate cyclases and phosphodiesterases that these bacteria encode (14). Accordingly, mutation of *pdeH*, expected to elevate c-di-GMP levels, has an inhibitory effect on swimming motility in both bacteria (10, 12, 15). A study that first demonstrated c-di-GMP-binding to the C-terminal PilZ domain of YcgR, also showed that the inhibitory effect of the *pdeH* mutant was abrogated by deleting *ycgR*, establishing later that YcgR::c-di-GMP is the only effector for motility inhibition (16, 17).

The flagellar motors of *E. coli* and *Salmonella* consist of a bi-directional rotor and dynamic stators that engage and disengage with the rotor in a load-dependent manner (18–22). The rotor includes the cytoplasmic C-ring made of three proteins (FliG, FliM, FliN). FliG is at the top of the ring, interlaced with inner membrane MS ring protein FliF, which in turn links to the periplasmic rod, external hook and the long helical flagellar filament. The stators are composed of pentameric MotA and dimeric MotB proteins (23, 24). They are independent units that drift in the inner membrane in a “closed” ion-nonconducting form until they encounter the flagellar basal body, upon which they dock on top of FliG in the C-ring and adopt an “open” ion-conducting conformation. Protons transiting through the MotAB complex cause conformational changes in MotA that power CW rotation of the pentameric MotA ring around a stationary MotB, thought to attach to peptidoglycan or some stationary moiety in the periplasm (25). Rotary MotA pushes on FliG, wherein a conserved patch of charged residues in MotA interact with compatible residues in the ‘torque helix’ of FliG, generating torque that results in the rotation of the C-ring and all parts linked to it (26). The C-ring can accommodate ∼11 stator units, but their numbers vary depending on external load. When attached to the flagellar filament (high load), *E. coli* motors spin at ∼125 Hz; when the filament is absent (low load), motors rotate at 300 Hz or faster (27, 28).

The default state of the bi-directional C-ring is CCW. The chemotaxis signal CheY∼P, interacts with FliM/FliN located below FliG to elicit a large change in the C-terminal domain of FliG (FliG_C_) where the torque helix is located (29), as well as producing changes at FliG_N_, concomitantly switching the orientation of charges in the torque helix of FliG and the rotational direction of the C-ring from CCW to CW (30–32). Between FliG_C_ and FliG_N_ is located a ‘cleft’ whose width changes from 30Å in CCW state to 40Å in the CW state. As first observed in *Borrelia burgdorferi* where the entire structure of the stator-associated rotor is visible, changes in FliG_C_ not only change the size of the FliG ring, but also change the disposition of the stators, which move inwards in the CW state and outward in the CCW state (33). Similar events at the C-ring were reported in *Salmonella* (30–32), where the diameter of the upper FliG ring toggles between ∼446 Å in the CW state to ∼466 Å in the CCW state (32). In the CW state, the rotating MotA pentamer would interact not only with FliG_C_ but also with FliG_N_, where charged residues similar to the torque helix in FliG_C_ are located. A genetic fusion between stators of a closely related bacterium and the torque helix in FliG_c_ (30), as well as docking simulations (31, 32), show that like seen in *B. burgdorferi*, stators must relocate in and out of the FliG ring.

YcgR action at the motor has been examined for over a decade, yet its mechanism remains unresolved. Some studies implicate the stators, others the rotor, and still others both. A persistent challenge is that YcgR interacts with multiple components—rotor proteins FliG and FliM and the stator protein MotA—with contacts mapped to the N- and C-termini of both FliG and YcgR. Recent rotor–stator cryo-EM structures revealing conformational changes during directional switching suggested a mechanism that we now explore experimentally.

## Results

### A new model for YcgR action

Multiple studies have demonstrated that in a *pdeH* mutant, where c-di-GMP levels are elevated, chromosomal YcgR produces two distinct motor phenotypes – increased CCW rotor bias and a reduction in motor speed (2–5, 7, 8, 13). In a *pdeH ycgR* double mutant carrying an inducible plasmid expressing YcgR, the rotor bias of some motors shifted to CCW before motor speed decreased (17), demonstrating that changes in bias and speed can occur sequentially. This temporal separation suggests that two mechanically distinct steps may be involved.

To explain these observations in the context of recent insights into the two C-ring conformations, we propose a model we call the **Texas 2-step**, delineated in Figure 1, which outlines the key components involved. The crystal structure of YcgR bound to c-di-GMP (6) is shown in Figure 1A. In this structure, the PilZ domain of YcgR (YcgR_C_) binds c-di-GMP between YcgR_N_ and YcgR_C_ (16), triggering a rearrangement of YcgR_C_ that exposes an α-helix at the extreme C-terminus (C-tail) (Fig. 1A, green helix) (6). The exposed C-tail of YcgR_C_ interacts with MotA residues previously identified as contacting the torque helix of FliG (26). This interaction is supported by mutations in both YcgR_C_ and MotA that abolish MotA-YcgR binding *in vitro* and YcgR-mediated motility inhibition *in vivo* (6). Consistent with this, extragenic suppressors of YcgR also map to the same MotA residues (2). Moreover, multiple studies have also demonstrated YcgR binding to FliG in the rotor (3, 4, 8).

**Fig 1.**
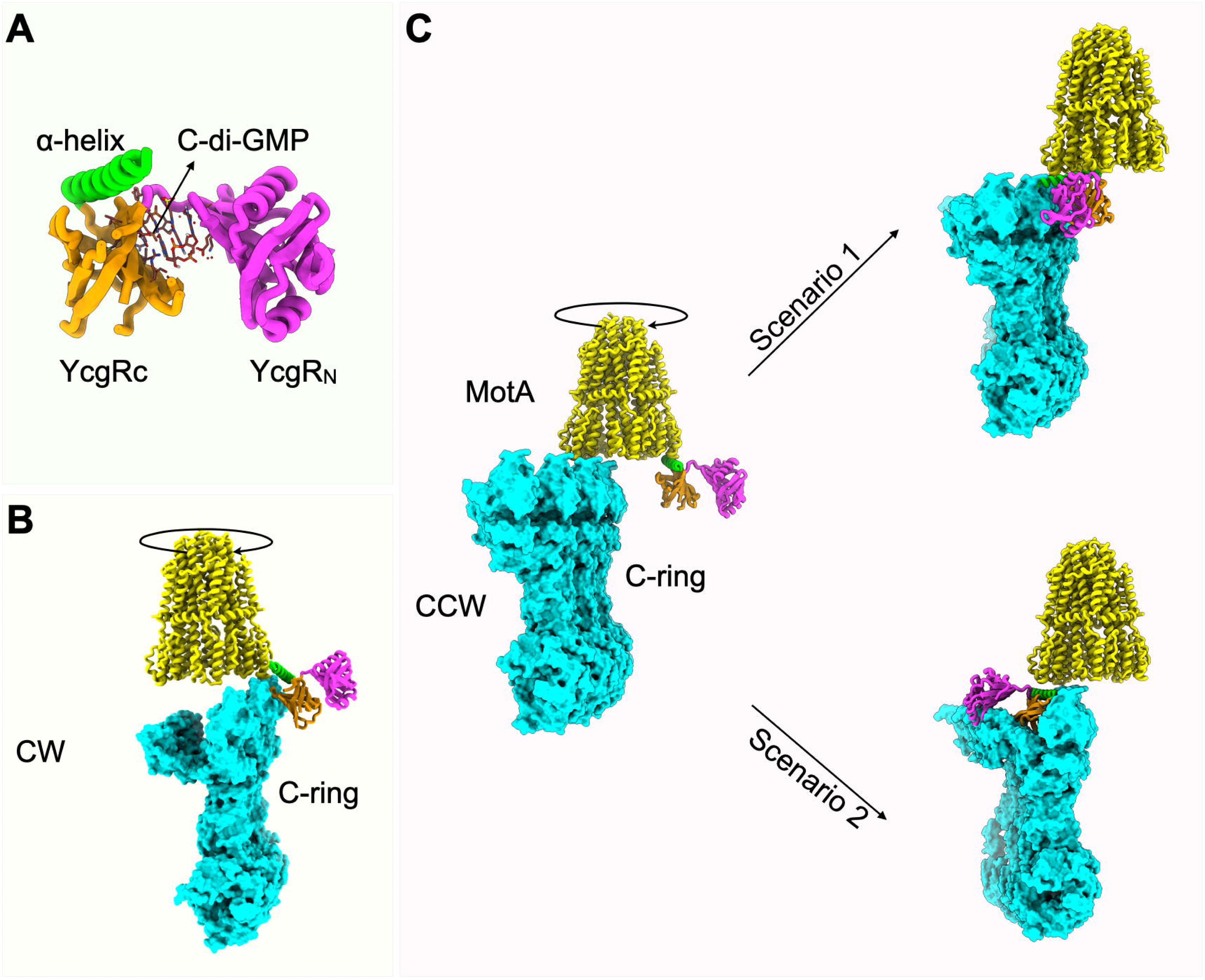
Two-Step Model of YcgR Action at the Motor. **A.** Crystal structure of YcgR bound to c-di-GMP (PDB: 5Y6F). The α-helix involved in MotA interaction is shown in green, C-terminal domain (YcgR_C_) in orange, and N-terminal domain (YcgR_N_) in magenta. **B.** CW (Clockwise) C-ring conformation (PDB: 8UMX; FliNMG subunits are in aqua) with a docked CW-rotating MotA pentamer (PDB: 6YKM; yellow). **C.** CCW (Counterclockwise) C-ring conformation (PDB: 8UMD). Unlike in B, the α-helix of YcgR has clear access to the bottom of MotA. This is Step 1. There are two possibilities for Step 2. In Scenario 1, as the MotA pentamer rotates, YcgR moves with it until YcgR_N_ contacts the C-terminal domain of a FliG subunit (FliG_C_). Straddling the MotA–FliG_C_ interface, YcgR obstructs MotA rotation. In an alternate Scenario 2, YcgR traverses the top of the C-ring to bind FliG_N_, settling into the cleft between FliG_N_ and FliG_C_.

### The model

In the first step, YcgRc can access MotA only in stators engaged with the CCW rotor conformation, where they project outward from the C-ring. This interaction is likely prevented in the CW conformation, where the stators are expected to shift inward, shielding the YcgR-binding region of MotA (compare Fig. 1B vs 1C).

In a second step, as the YcgR_C_-bound MotA pentamer begins to rotate, YcgR_N_ is positioned to contact FliG. We envision two possible scenarios for this interaction. In the first, YcgR bridges FliG_C_ and MotA via its two domains, either arresting rotation of the bound MotA pentamer (Fig. 1C, Scenario 1; Movie 1), or more plausibly, remaining attached to FliG_C_ while MotA dissociates (not shown). In the second scenario, YcgR disengages from rotating MotA and moves across the top of the C-ring to bind FliG_N_ (Fig. 1C, Scenario 2; Movie 2) from where it may weaken or prevent MotA binding. In both cases, YcgR occupation of FliG stabilizes the CCW state, reducing rotation speed as additional YcgR molecules engage FliG.

Involvement of FliG in our proposed second step is supported by several lines of evidence. Genetic analyses show that mutation in the FliG_C_ torque helix residues relieve YcgR-mediated motility inhibition (4). Two-hybrid and pull-down assays demonstrate that YcgR interacts with both FliG_N_ and FliG_C_ (3). Moreover, TIRF experiments measuring eGFP-YcgR fluorescence at the motor before and after addition of NaN_3_ – an uncoupler that dissipates the proton motive force and causes stators to dissociate – showed no change in fluorescence intensity, indicating that YcgR remains stably associated with the rotor (5). The closest rotor protein to MotA-bound YcgR is FliG, consistent with these observations.

Involvement of YcgR_N_ in motility inhibition is also supported by genetic evidence: even when the MotA-interacting α-helix in YcgRL remains intact and functional, mutations in YcgR_N_ residues disrupt motility (4, 8). Thus, interaction of YcgR with MotA alone cannot fully account for all YcgR-dependent effects.

In this study, we focus exclusively on **Step 1** of the proposed model, which makes a clear prediction: this step should be blocked in CW-locked rotors.

### Testing the first step of the model

We used two *E. coli* parent strains - HCB5 and MG1655 - to ensure that observed motor behaviors were not strain-dependent. HCB5 carried only a *pdeH* deletion (WT*), while MG1655 was deleted for both *pdeH* and *ycgR* (WT**). To generate motors that rotated predominantly in one direction, we introduced into these strains deletions in genes encoding the chemotaxis signaling components or the rotor protein FliG: Δ*cheY* - CCW, Δ*cheZ* - CW, *fliG*_ΔPAA_ (deletion of PAA residues 169-171) - extreme or ‘locked’ CW (34). In both strain backgrounds, YcgR was supplied from an inducible plasmid, as described previously (17).

Motor behavior was monitored using a bead assay, where a 0.75 μm polystyrene bead was attached to a stub of a sheared flagellar filament on bacteria fixed to a glass slide, and bead rotation was captured by high-speed videos (see Methods). For most strains, at least 15 motors were recorded continuously for 10 min following inducer addition.

The behavior of the *pdeH ycgR* (WT**) strain upon YcgR induction was reported previously (17) and is reproduced here in Fig. S1A (top trace). The *pdeH* strain (WT*) showed a similar response (Fig. 2A and S1A bottom trace). This strain expresses chromosomal levels of YcgR, and the motors display the expected variation in CCW bias and speed. Upon plasmid-driven YcgR induction, we observed three distinct motor behaviors: a strong CCW bias with no reduction in initial speed (Fig. 2A, top trace); a CCW bias accompanied by a gradual speed reduction (Fig 2A, bottom trace); or no change in bias prior to sudden motor arrest (Fig. S1A, bottom trace). These observations indicate that the motor response to YcgR induction is independent of the strain background.

**Fig 2.**
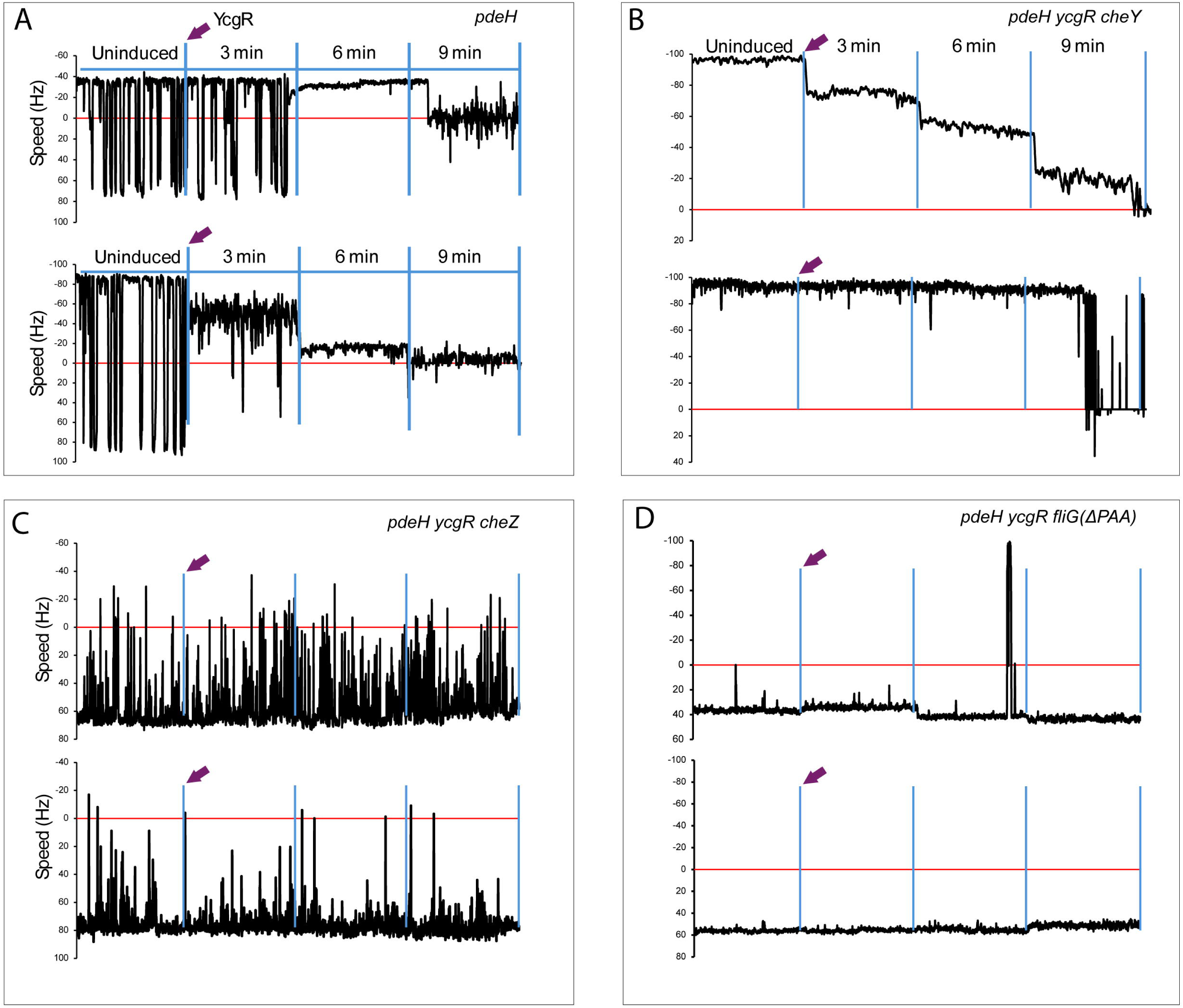
Response of WT, CCW and CW motors to YcgR. **A.** Two representative motors from WT* HCB5 strain are shown. YcgR was supplied from an IPTG-inducible low copy plasmid pSEVA224. Motor behavior was monitored by recording the motion of 0.75 μm polystyrene beads attached to flagella filament stubs (see Methods). After IPTG addition (arrow), 60-second representative segments from each consecutive 3-minute recording interval are shown. CCW (-) and CW (+) speeds in Hz are shown on the Y-axis. The profiles are representative of 15 individual motors tracked (see Fig. 3). **B.** As in A, except the strain is WT** MG1655 and has *cheY* deleted. **C.** As in B, except *cheZ* is deleted. **D.** As in B, except *fliG* is deleted for nucleotides encoding PAA at residues 169-171. Relevant genotypes of the strains are indicated on the top right-hand corner.

The Δ*cheY* strain, which has an initial CCW bias, also displayed two behaviors: a graded speed reduction over time (Fig. 2B, top trace), or an abrupt stop without prior speed reduction (Fig. 2B, bottom trace).

The two CW-biased strains - Δ*cheZ* and FliGΔPAA – were largely impervious to YcgR (Fig. 2C-D). Δ*cheZ* motors rotated predominantly CW, with an ‘unstable’ bias characterized by occasional excursions into the CCW zone, yet they remained mostly unresponsive to YcgR (Fig. 2C). When a larger bead (0.99 μm) was used, the CW bias of Δ*cheZ* motors became even more unstable, switching rapidly between the two conformations (Fig. S1B, traces before YcgR induction). Upon YcgR induction, two Δ*cheZ* motors exhibited a shift to CCW accompanied by speed reduction (Fig. S1B, top and bottom traces at 9 and 6 min, respectively), followed in one case by a pronounced CCW bias and speed reduction (Fig. S1B, bottom), resembling the motor traces shown in Figure 3A. These observations indicated that even Δ*cheZ* motors that respond to YcgR do so primarily in the CCW regime.

**Fig 3.**
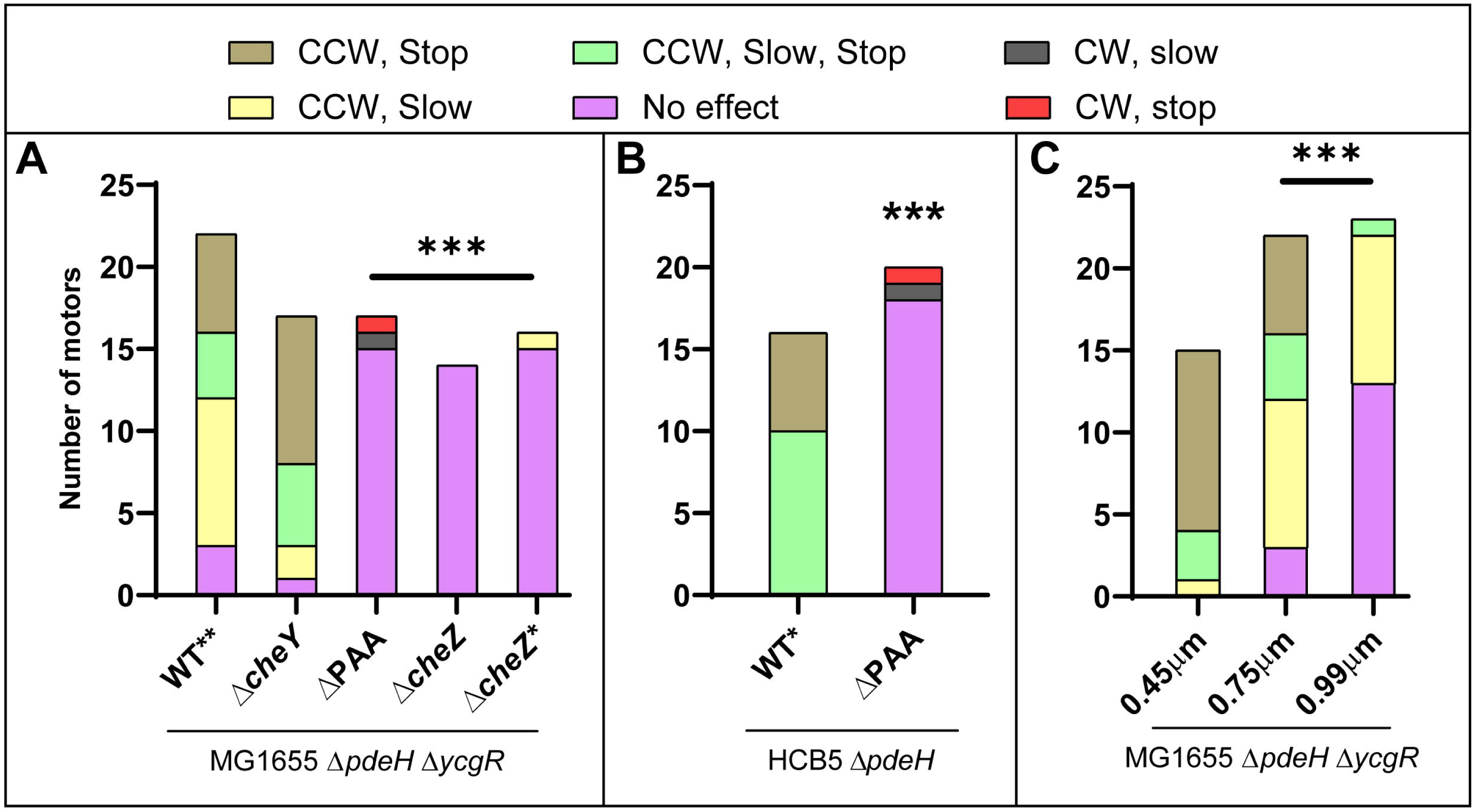
Summary of all motor traces. **A,B.** The relevant mutation expected to affect motor behavior is indicated directly under each bar graph. Genotype of WT* and WT** strains, common to all motors, is under the line. Bead size is 0.75 μm for all graphs, except for Δ*cheZ*\*, where the size was 0.99 μm. Color key indicates the motor behaviors observed. **C.** Behavior of WT** motors under varying bead sizes. p-values were calculated only for motors showing ‘no effect’ compared to WT. Fisher’s exact test was used to assess statistical significance; *** indicates p < 0.001.

FliGΔPAA motors displayed the expected stable or ‘locked’ CW bias, with most motors resistant to YcgR (Fig. 2D, top and bottom). However, in both WT* and WT**, approximately 10% of the CW-locked motors showed either a gradual decrease in speed (Fig. S1CD, top), a gradual decrease followed by a stop (Fig. S1C, bottom), or a sudden stop (Fig. S1D, bottom). These observations indicate that YcgR can access a minority of CW-locked motors. We note that the cleft between FliG_N_ and FliG_C_ is larger in the CW rotor, and that a density consistent with YcgR has been observed in one data set from FliGΔPAA motors (31).

A summary of the motors tracked in this study is shown in Figure 3. While majority of the data were generated with 0.75μm beads, we also monitored WT** motors with smaller (0.45μm) and larger (0.99μm) beads, which are expected to impose lower and higher loads on the motor, respectively (Fig. 3C). A clear trend emerged: upon YcgR induction, more motors stopped when smaller beads were used compared with larger ones. This trend is consistent with the expected relationship between bead size and motor torque, as smaller beads engage fewer stators at the rotor, while larger beads engage more, explaining the load-dependent differences in motor response.

## Discussion

Using strains whose motors rotate predominantly in the CCW or CW directions, we show that YcgR primarily affects motors in the CCW conformation, supporting the proposed first step our model (Fig. 1C). Dependence on the CCW conformation would facilitate YcgR access to the stators, where YcgR binds MotA—an interaction supported by genetic and biochemical evidence (2, 6). Binding of YcgR to MotA is expected to disrupt MotA’s normal torque-generating interaction with FliG, thereby reducing motor speed. This first step, however, cannot fully account for YcgR function, as YcgR also alters rotor bias.

We therefore propose a second step in which YcgR_N_ binds FliG_C_, the rotor subunit most proximal to MotA-bound YcgR (Fig.1, Scenario 1) (see ‘A new model’ above for supporting evidence). Such an interaction would drive the rotor towards a CCW conformation, coupling bias stabilization with progressive speed reduction, and eventually leading to motor arrest as more YcgR molecules engage the rotor (Movie1).

Scenario 1 in the proposed second step could, in principle, explain both the CCW bias shift and the accompanying speed reduction observed in WT motors. To account for WT motors that shift to a CCW bias **without** an initial decrease in speed (Fig. 2A & S1A, top traces), one would have to assume that YcgR can remain bound to the rotating MotA pentamer without affecting torque, which seems unlikely. Instead, Scenario 2 - where YcgR_N_ binds FliG_N_ - better accommodates this behavior. This scenario is supported by findings that YcgR interacts with both FliG_N_ and FliG_C_ in two-hybrid and pull-down assays (3). A trivial explanation for these dual interactions could be that sequence homologies between the two FliG domains reflect their shared MotA-contacting surfaces in the distinct rotor conformations. Figure 4 presents a final model, shown as an aerial view of the C-ring, summarizing support for Step 1 from the data obtained in this study.

**Fig 4.**
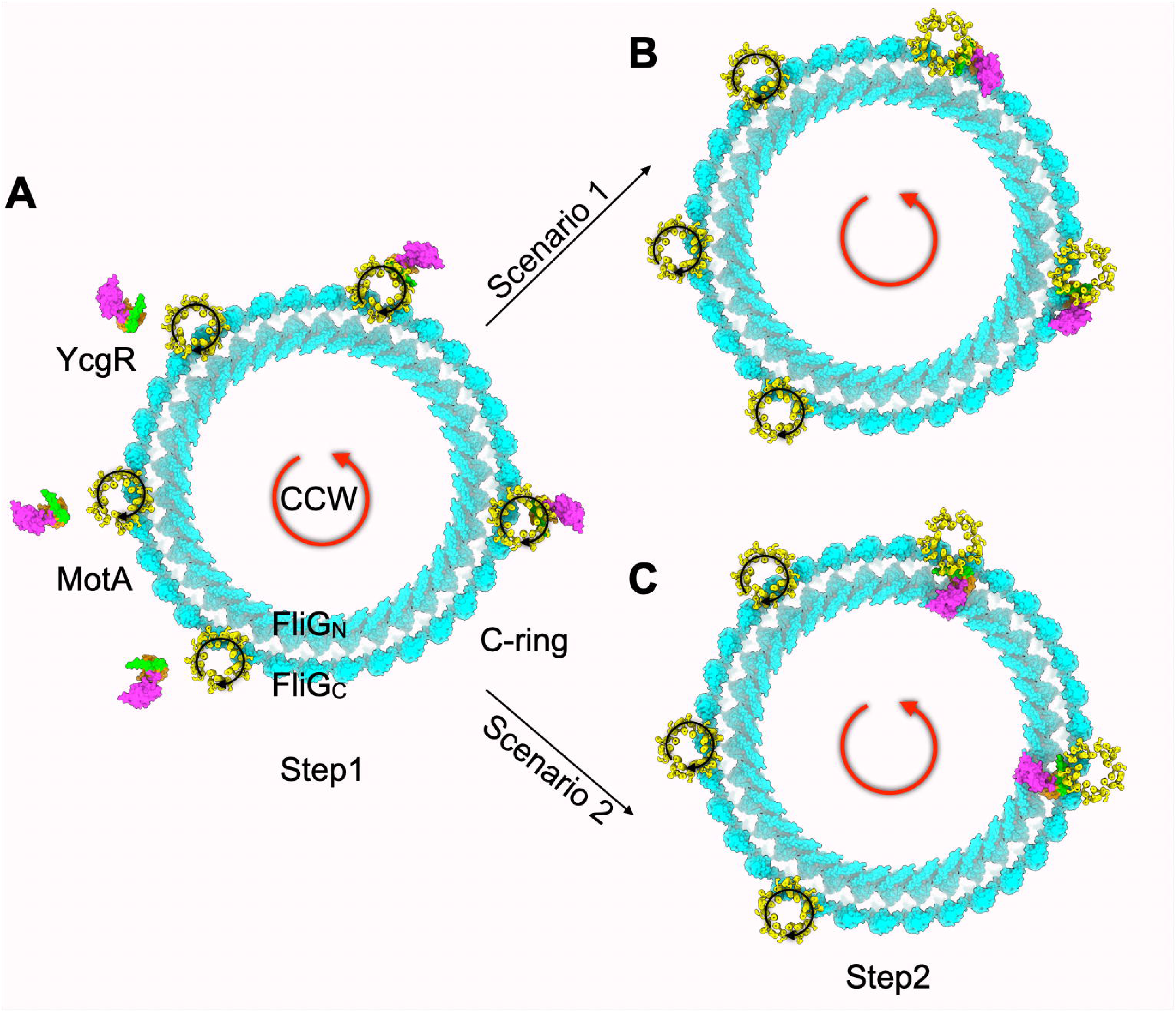
Final model with an aerial view of the FliG ring. **A.** YcgR only affects motors in the CCW conformation, supporting the proposed first step of our model. The MotA pentamer is accessible for binding in this conformation. **B.** To affect both speed and bias, YcgR must engage with both the stator and the rotor. This second step could be achieved by delivery of YcgR to either the outer (FliG_C_) or inner (FliG_N_) FliG rings by the rotating MotA pentamer.

### Additional Mechanistic Insights

1. Given the variety of experimental protocols demonstrating binding of YcgR to the rotor (3–5, 8), why did extragenic suppressors of YcgR only map to MotA residues that interact with FliG (2), and not to FliG itself? To address this question, we re-isolated YcgR suppressors and recovered the same MotA mutations - G93E, G93R, G93V and S96L – along with a new mutation, Q91P. None mapped to *fliG*. We also obtained these suppressors from the Jenal lab (Basel, Switzerland) and characterized their motor behavior (Fig. S2). The suppressors were identified by their ability to swim in soft agar plates (Fig. S2A), an assay that requires functional chemotaxis. Motor assays revealed that the suppressors exhibited normal motor bias but reduced rotation speeds (50–70 Hz) compared to wild type (80 Hz) (Fig.S2 B,C). These reduced speeds were intrinsic to the suppressor mutations, as they persisted even in the absence of YcgR (Fig. S2B, Δ*ycgR,* S96L shown). We consider two explanations for why YcgR suppressors did not map to *fliG*. First, because YcgR can interact with both FliG_N_ and FliG_C_, the probability of mutationally disrupting both binding sites is low. Alternatively, mutations in these regions of FliG may compromise motor bias and thus fail to be recovered in the chemotaxis-dependent motility screens used for suppressor isolation. Indeed, mutations targeting the torque helix of FliG severely impair motility in soft agar (4).
2. In a FRAP study using TIRF microscopy, 36% of mEGFP-YcgR molecules expressed in a *pdeH* mutant were found to be weakly bound to the motor, with the remaining fraction tightly bound (7). Evidence that the tightly bound YcgR population localizes to the rotor rather than the stator comes from a similar TIRF-based analysis showing that depletion of the proton motive force did not alter YcgR fluorescence (5). Moreover, two independent studies that directly interrogated interactions with both MotA and FliG identified MotA as the primary target of YcgR (6, 8). Taken together, these findings suggest that YcgR initially associates with MotA, which induces a conformational change that enables its subsequent tight binding to FliG.
3. Motor behavior observed at chromosomal levels of YcgR (Fig. S2; *pdeH*) as well as at plasmid-induced levels (Fig. S1B; *pdeH ycgR cheZ*) shows that, despite reduced speed and a CCW bias, these motors are still capable of switching. We envision two possibilities: (i) YcgR-bound FliG subunits do not impede rotor switching, or (ii) YcgR-occupied subunits become functionally or spatially isolated from the remainder of the rotor, which continues to switch normally.
4. Significant binding of YcgR to FliM has been reported in only one study, which employed three independent approaches—pulldown assays, visualization of YcgR–GFP at the rotor, and motility analyses (4). Interestingly, two YcgR variants, R118D and S147A, which differ in their ability to bind c-di-GMP and modulate motor function, were both found to interact with FliM, albeit more weakly than with FliG (3, 4, 16). These findings collectively make a credible case for YcgR–FliM interaction. Given that the R118D mutant, reported to be defective in c-di-GMP binding, was nevertheless observed at the motor (4), we consider the possibility that FliM serves as a reservoir for YcgR, positioning it to engage the motor rapidly upon an increase in intracellular c-di-GMP levels.
5. In contrast to *E. coli* YcgR, the *Bacillus subtilis* homolog MotI (also known as DgrA) does not influence motor bias but instead functions as a molecular clutch, binding to MotA and disengaging it from the C-ring to halt rotation (35). The evolutionary advantages of employing distinct mechanisms to control motor activity through these related proteins remain speculative. Nevertheless, their behaviors converge under certain conditions: at subinhibitory concentrations, MotI mimics YcgR by reducing swimming speed, whereas at high concentrations, YcgR mimics MotI by stopping the motor.

## Acknowledgments

This model was inspired by the first published images of the *Borrelia* motor from Jun Liu’s lab showing differences in stator disposition at the rotor; we thank him for discussions. We also thank Urs Jenal and Birgit Scharf for strains. This work was supported by NIH grant GM118085 to RMH.

## Methods

### Strains and growth conditions

Strains and plasmids used in this study are listed in Table S1. Bacterial cultures were grown in Lennox broth (LB) base (20 g/liter) (Fisher BioReagents). For motility assays in soft agar, 8 μl of an exponential-phase culture (optical density at 600 nm [OD_600_], ∼0.6) was inoculated onto plates made with 0.3% Bacto agar (Difco) and incubated at 30°C. All plate images shown are representative of three biological replicates, each in triplicate. Where required, the following antibiotics were used: ampicillin (100 μg/ml), chloramphenicol (20 μg/ml), and kanamycin (50 μg/ml). For inducible plasmids, 50uM IPTG (isopropyl-β-d-thiogalactopyranoside) and 0.2% L-arabinose were used.

### Genetic manipulation

The WT parent strains for *E. coli* were MG1655 and HCB5. Mutant strains were constructed by inserting a kanamycin resistance cassette (KAN) into the designated gene as previously described (36) or sourced from the Keio collection (37). Mutations were transferred to fresh backgrounds by phage P1 (P1 Cm) transduction. Excision of the inserted KAN cassettes was achieved by expression of the FLP recombinase encoded by pCP20 (36).The resulting strains were confirmed by DNA sequencing. A 3 amino acid deletion of residues 169-171 (PAA) in FliG was introduced using the CRMAGE protocol (38), which leverages CRISPR-Cas9 technology as follows: A cocktail of 3 primer pairs for making guide RNAs (gRNA) targeting *fliG* were cloned into pMAZ-SK plasmid via Golden Gate assembly; gRNAs are expressed with anhydrotetracycline (aTetracycline). These constructs were introduced along with repair oligonucleotides into NBN28 containing pMA7CR_2.0, which expresses the λ Red β-protein and the CRISPR/Cas9 protein upon induction with L-arabinose. The insertion was confirmed by PCR amplification followed sequencing. All the primers and oligonucleotides used are listed in Table S2.

### Measurement of motor dynamics using bead assay

The bead assay was conducted as described in the previous work (17, 39). *E. coli fliC* deletion strains (containing additional chromosomal mutations) were complemented with pFD313 expressing the ’sticky’ FliC variant (40), and pSEVA224 expressing YcgR. First, cells were passed through two syringes connected by a 7” polyethylene capillary to shear the flagella. A 40 µL cell suspension was then added to a poly-l-lysine-coated cover slip, incubated for 10 minutes, and rinsed with motility buffer to remove unattached cells (41). A 40 µL solution of polystyrene beads (1:50 dilution) was added and left for 10 minutes before excess beads were washed away, prior to introducing IPTG for inducing YgcR expression. Bead rotations were captured at 1,000 frames/s using a CCD camera (ICL-B0620M-KC0; Imperx, Boca Raton, FL). Videos were processed using custom analytical programs within LabVIEW 2012 as described before (17).

### Animation

Animation illustrating YcgR-mediated regulation of the stator–C-ring complex. was created using ChimeraX 1.9 and assembled in Keynote. Structural models used include YcgR (PDB: 5Y6F), C-ring in CW and CCW conformations (PDB: 8UMX and 8UMD, respectively), and MotA (PDB: 6YKM).

