## Supplementary Figures for "Texas 2-Step: A new Model for YcgR::c-di-GMP Action at the Flagellar Motor"

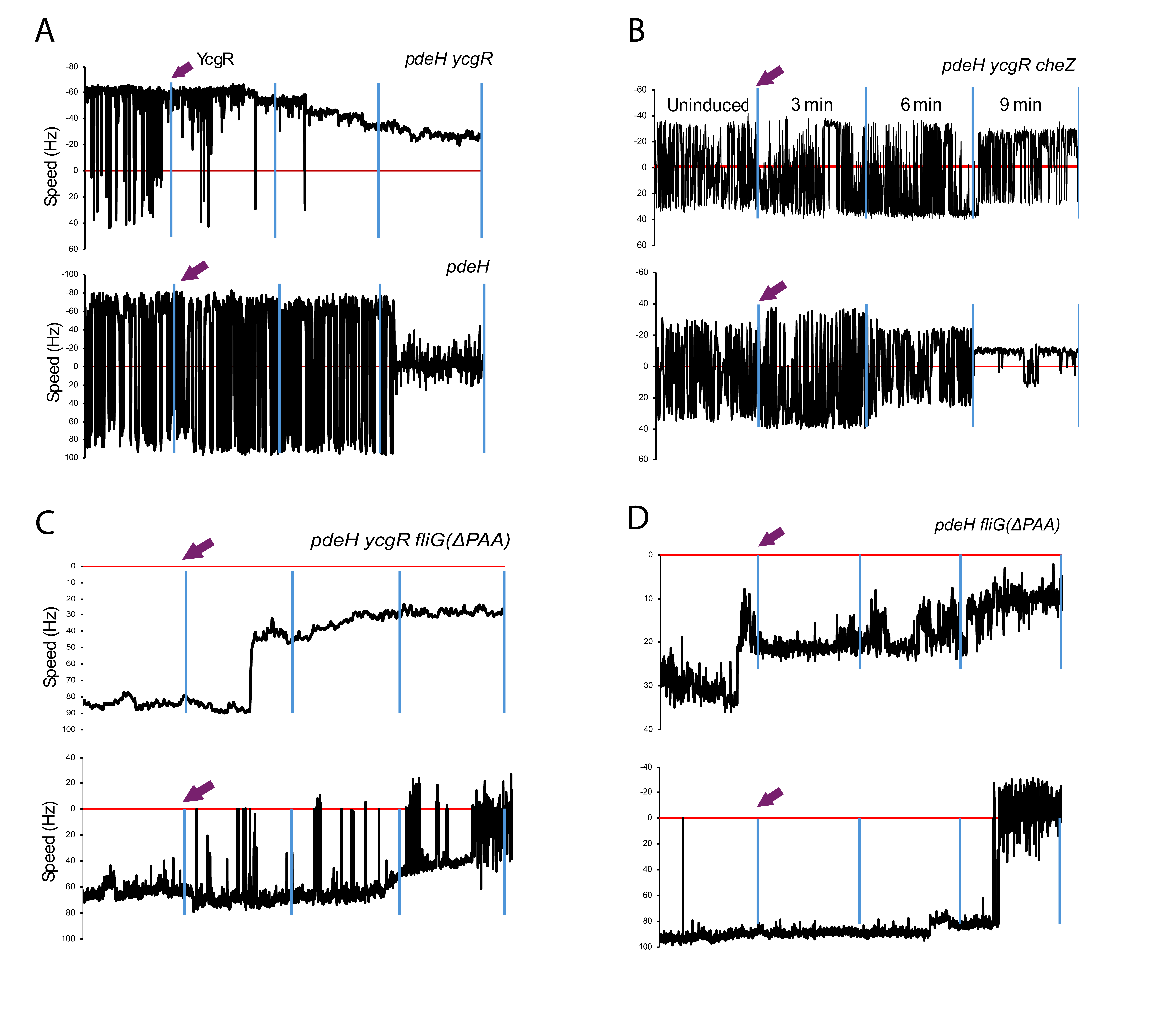
**Supplementary Figures and Movie Legends**

**Fig S1.** Behavior of motors in varied backgrounds and bead sizes. **A.** Two representative motors from WT** MG1655 strain. All other descriptions as in Fig 2A. **B**. Behavior of two *cheZ* motors when the bead size was 0.99 μm. **C,D.** Representative *fliG* (ΔPAA) motors in WT* and WT** backgrounds. Bead size was 0.75 μm. Other descriptions as in Fig. 3A.

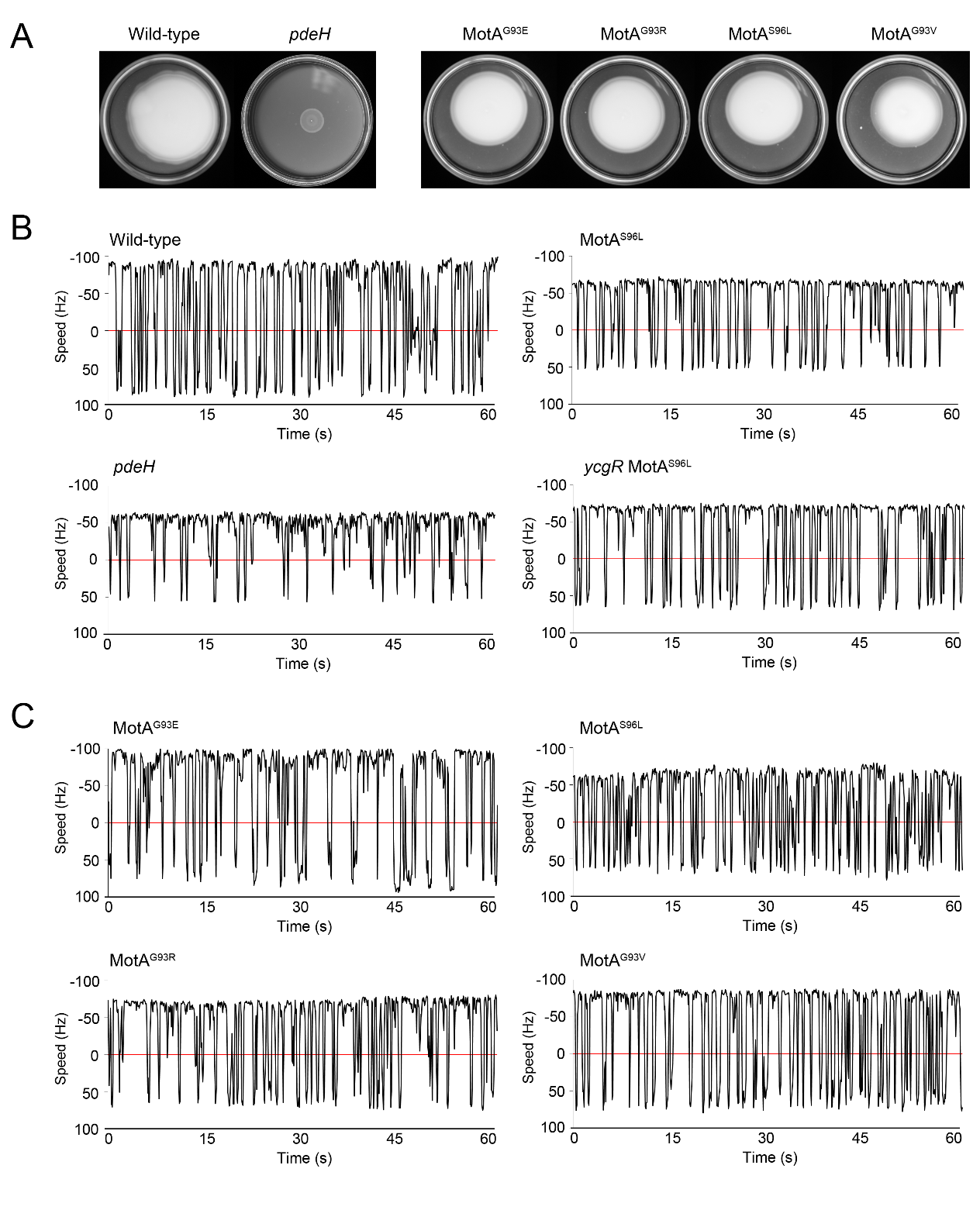

**Fig. S2**. Swimming and motor behavior of MotA suppressors. **A.** Motility assays were conducted in soft agar (see Methods). Left panel, motility is inhibited in a *pdeH* mutant (WT, MG1655; *pdeH*, AB607). Right panel, four MotA suppressors that overcame the motility defect of the *pdeH* mutant, described by Boehm et al. (see Table S1 for strain numbers). **B.** Bead assays in WT and indicated MotAS96L suppressor (AB1577) in which *ycgR* was deleted (JP1501). **C.** Bead assays in four MotA suppressor strains. The WT strain is MG1655 for all motors, monitored with 0.75 μm polystyrene beads. Representative traces of 15 motors each are shown.

**Movie 1**. YcgR interaction with CCW stator–C-ring complex, with the two YcgR domains straddling MotA and FliG_C_. The animation was created using ChimeraX 1.9 and assembled in Keynote. Structural models used include YcgR (PDB: 5Y6F), CCW C-ring (PDB: 8UMD), and MotA (PDB: 6YKM).

**Movie 2.** YcgR interaction with CCW stator–C-ring complex, with YcgR binding FliG_N,_ settling into the cleft between FliG_N_ and FliG_C_. Structural models as in Movie 1.
