## Supplementary Tables for "Texas 2-Step: A new Model for YcgR::c-di-GMP Action at the Flagellar Motor"

**Table S1. Strains and Plasmids**

| **Strain** | **Genotype/Description** | **Source/reference** |
| --- | --- | --- |
| MG1655 | Wild type *E. coli* F^-^ λ^-^ *ilvG*- *rfb*-50 *rph*-1 | Lab collection |
| NBN28 | MG1655 ∆*fliC* | This work |
| NBN77 | MG1655 ∆*pdeH* ∆*ycgR* | This work |
| NBN44 | NBN77 ∆*fliC* | This work |
| NBN43 | NBN44 ∆*cheZ* | This work |
| NBN68 | NBN44 ∆*cheY* | This work |
| NBN65 | NBN28 *fliG* (∆169-171) | This work |
| NBN89 | NBN65 ∆*pdeH* ∆*ycgR* | This work |
| HCB5 | AW405 ∆*fliC* | Scharf Lab, (1) |
| NBN116 | HCB5 ∆*pdeH* | This work |
| NBN121 | NBN116 *fliG* (∆169-171) | This work |
| AB434 | MG1655 ∆*ycgR::Frt* | (2) |
| AB607 | MG1655 ∆*pdeH::Frt* | (2) |
| AB1468 | AB607 1976787::Tn*mariner*(kan) *motA*-1 (G93E) | (2) |
| AB1576 | AB607 1976787::Tn*mariner*(kan) *motA*-4 (G93R) | (2) |
| AB1577 | AB607 1976787::Tn*mariner*(kan) *motA*-2 (S96L) | (2) |
| AB1578 | AB607 1976787::Tn*mariner*(kan) *motA*-3 (G93V) | (2) |
| JP1501 | AB1577 ∆*ycgR* | This work |

| **Plasmid** | **Expressed Protein** | **Host Plasmid** | **Resistance** | **Induction** | **Reference** |
| --- | --- | --- | --- | --- | --- |
| pCP20 | FLP recombinase | **N.A*** | Ampicillin | Constitutive | (3) |
| pSEVA224 | YcgR | N.A. | Kanamycin | IPTG | (4) |
| pFD313 | FliC^sticky^ | pTRC99a | Ampicillin | IPTG | (5) |
| pMA7CR_2.0 | λ Red β-protein and Cas9 | pMA7 | Ampicillin | L-arabinose and **aTc*** | (6) |
| pMAZ-SK | gRNA | pCOLA-duet | Kanamycin | aTc | (6) |
| pBAD24 | Cloning vector | N.A | Ampicillin | L-arabinose | (7) |
| pVN8 | YcgR-GFP | pBAD24 | Ampicillin | L-arabinose | (8) |

**N.A***: Not applicable

**aTc***: Anhydrotetracycline

**Table S2. Primers pairs for gRNA and repair oligonucleotide for FliG deletion**

| **Name** | **Sequence** |
| --- | --- |
| 1_Fwd | TAGTGGCTGGCTGCACGCCGCCAA |
| 1_Rev | AAACTTGGCGGCGTGCAGCCAGCC |
| 2_Fwd | TAGTGTCAGCTCCGCCAGCGCGGC |
| 2_Rev | AAACGCCGCGCTGGCGGAGCTGAC |
| 3_Fwd | TAGTGAGCAAGCCATTCAGTACTT |
| 3_Rev | AAACAAGTACTGAATGGCTTGCTC |
| *fliG*_Repair oligos | CTGCGCCACGACGTGATGTTGCGTATCGCCACATTTGGCGGCGTGCAGCTGGCGGAGCTGACAGAAGTACTGAATGGCTTGCTCGACGGTCAGAATC |
